# Evaluation of Osteoinductivity of Different Calcium Phosphate and PTMC-Calcium Phosphate Composite Biomaterials in a Sheep Model

**DOI:** 10.1101/666800

**Authors:** Arden L. van Arnhem, Ni Zeng, Anne van Leeuwen, Roel Kuijer, Ruud. R.M. Bos, Dirk W. Grijpma

## Abstract

Osteoinduction refers to *de novo* bone formation induced by biomaterials in places where physiologically no bone tissue is formed. Biomaterials with osteoinductive capacities have been shown to fill bone defects of critical sizes with ubiquitous new bone formation. Therefore, osteoinduction has been regarded as an important characteristic for biomaterials aiming at bone regeneration. In our study, we tested osteoinductive capacities of different calcium phosphate bioceramic particles, calcium phosphate scaffolds, and porous poly(trimethylene carbonate)(PTMC)-calcium phosphate composite scaffolds in a sheep model. Biphasic calcium phosphate (BCP) particles of 45-150 μm and 150-500 μm, microporous β-tricalcium phosphate (β-TCP) particles of 45-150 μm, non-microporous β-TCP particles of 45-150 μm and 150-500 μm, and porous β-TCP scaffolds were implanted in sheep long dorsal muscle for three and nine months. Likewise, porous composite scaffolds, in which BCP particles, microporous β-TCP particles and β-TCP particles, all of 45-150 μm, had been incorporated into PTMC matrices, were implanted in sheep long dorsal muscle for three and nine months. Porous PTMC scaffolds were implanted as controls. Abundant new bone formation was induced by BCP particles of both size ranges, the β-TCP scaffold was also able to induce new bone formation at both three and nine months follow up, while no new bone formation was induced by the other biomaterials. Implantation of the abovementioned biomaterials led to uneventful degradation of the PTMC matrices and the incorporated calcium phosphate particles, and provoked no obvious tissue reaction. Future studies are needed to determine the optimal composition of composite biomaterials based on PTMC and calcium phosphate to produce osteoinductive composites.

## INTRODUCTION

New bone formation in physiological remodeling and repairing of damaged bone tissue occurs via osteogenesis, which involves recruiting mesenchymal stem cells, mainly from mesodermal tissues such as bone marrow, and inducing their proliferation and differentiation to osteoblasts at the sites(1). Concerning reconstructions of bone defects by biomaterials, the term ‘osteoconduction’ is used to describe new bone formation stimulated and guided on surfaces or into pores provided by the biomaterials implanted in bone defects. Besides, a phenomenon called ‘osteoinduction’, defined as “the induction of undifferentiated inducible osteoprogenitor cells that are not yet committed to the osteogenic lineage to form osteoprogenitor cells”(2), describes induced new bone formation in the biomaterials implanted in ectopic sites, such as subcutaneous or intramuscular implantations(3). Biomaterials with osteoconductive properties ‘promote the recruitment and migration of osteogenic cells into the wound site’(4) and serve as scaffolds for new bone formation to occur, while biomaterials with osteoinductive properties actively induce new bone formation and are believed to have the essential ability to heal bone defects of critical sizes successfully(5). Compared to what occurs in osteoconductive biomaterials, new bone formation induced by osteoinductive biomaterials occurs not only at the interface between the biomaterials and the host tissue, but also all over the defects filled with the biomaterials(6). Thus, it seems attractive and beneficial to apply osteoinductive biomaterials to reconstruct bone defects of critical sizes, since new bone formation led by osteoconductive biomaterials cannot fill up most such bone defects. Although still not fully understood, mechanisms of osteoinduction have been extrapolated to include a direct induction of recruitment, proliferation and differentiation of mesenchymal stem cells from blood, fat or muscle tissue to cells in the bone forming lineage by osteoinductive biomaterials, and an indirect induction by proteins which induce new bone formation and are absorbed to the osteoinductive biomaterials during *in vivo* implantations(1, 7). Autologous bone grafts contain viable precursor cells for osteogenesis and possess excellent biological and mechanical properties for both osteoconduction and osteoinduction, thus they have been regarded as the ‘golden standard’ in reconstructing bone defects in oral and maxillofacial surgery(8). Drawbacks in using autologous bone grafts to reconstruct bone defects include limited availability of donor sites(9), morbidity in donor sites, risks of infections, nerve damages, hemorrhage, prolonged surgical procedures(10, 11) and unpredictable resorption of autologous bone grafts after implantation(12–14). These drawbacks impose necessities of using and developing bone graft substitutes with similar biological and mechanical properties as well as clinical performances. Among different bone graft substitutes, synthetic biomaterials are of especially high research interest, because they can be designed to possess bioactivities similar to autologous bone grafts and can be produced in controlled manners and in large amounts. Synthetic calcium phosphate bioceramics, a prominent class of bone graft substitutes, have been widely used in trauma and orthopedic surgery in the Netherlands(15), and are considered as good alternatives for autologous bone grafts(15, 16) because they provide a source of calcium ions and phosphate ions, which are necessary for new bone formation in bone defects, during their degradation. Hydroxyapatite (HA), which appears as the inorganic component in natural bone tissue, provides good mechanical support in bone defects in non-load bearing sites, has been used as bone graft substitutes, it takes years before a full degradation(17). HA bioceramics are reported to show osteoinductive potential in dogs(18, 19) and baboons(20) when they are implanted subcutaneously or intramuscularly. β-tricalcium phosphate (β-TCP) bioceramics degrade faster than HA bioceramics due to their higher solubility *in vivo*, possess good osteoconductive capacity and serve as excellent scaffolds for bone regeneration in bone defects(21, 22). Biphasic calcium phosphate (BCP) bioceramics contain HA and β-TCP crystalline structures at different ratios, often 60% of HA with 40% β-TCP(23) or 80% HA with 20% β-TCP(24). BCP bioceramics in forms of particles, blocks or injectable substances turn to be highly promising bone substitutes for uses in orthopedic, dental and maxillofacial surgeries thanks to their good bioactivities derived from the combination of HA and β-TCP(25–27). Although chemical compositions of different calcium phosphate bioceramics are similar or even identical, their bioactivities can significantly differ due to differences in porosities in macro- and micro-view(15), sintering temperature(24), the ratio of calcium and phosphorus(28), particles sizes(29), overall geometry(30), surface roughness and specific surface area(31). Currently it is still to be determined what parameters make calcium phosphate bioceramics fully resemble autologous bone grafts and subsequently replace their use.

Despite excellent bioactivities and biological performances of calcium phosphate bioceramics *in vitro* and *in vivo*, their inherent high brittleness prevents their plastic deformation and hinders their wide clinical application, especially in load bearing sites (17, 32). A feasible solution to such a problem is to incorporate calcium phosphate bioceramics into polymeric matrices, since natural bone tissue is essentially mineralized collagen matrices(33).

The presented study first aims to test osteoinductive properties of six different calcium phosphate bioceramics, with a special interest in the influence of granule sizes in osteoinduction, in a sheep model. Since BCP particles in the size range of 45-150 μm have been shown to be osteoinductive(34), the other aim of our study is to investigate whether incorporation of BCP particles and two other β-TCP particles in the same size range into PTMC matrices makes the composite scaffolds osteoinductive as well. Besides, the biodegradation and biocompatibility of all tested calcium phosphate biomaterials and PTMC-calcium phosphate composite scaffolds are studied.

## MATERIALS AND METHODS

### Materials

Table 1 presents the physiochemical and structural characteristics of the different calcium phosphate and PTMC-calcium phosphate composite biomaterials included in our study.

**Table 1.**
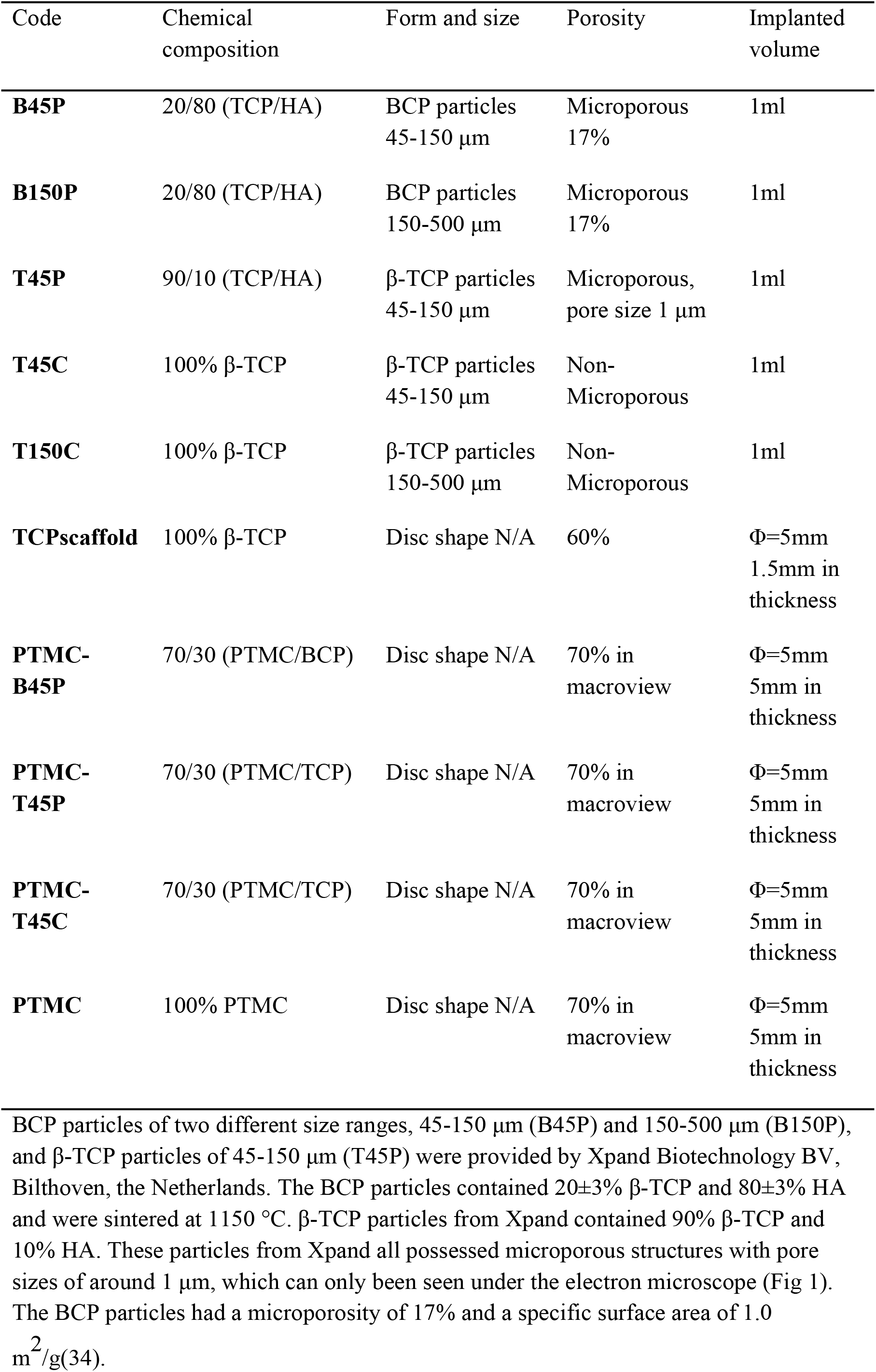
Overview of the included biomaterials

| Code | Chemical composition | Form and size | Porosity | Implanted volume |
| --- | --- | --- | --- | --- |
| **B45P** | 20/80 (TCP/HA) | BCP particles<br>45-150 $\mu\text{m}$ | Microporous<br>17% | 1ml |
| **B150P** | 20/80 (TCP/HA) | BCP particles<br>150-500 $\mu\text{m}$ | Microporous<br>17% | 1ml |
| **T45P** | 90/10 (TCP/HA) | β-TCP particles<br>45-150 $\mu\text{m}$ | Microporous,<br>pore size 1 $\mu\text{m}$ | 1ml |
| **T45C** | 100% β-TCP | β-TCP particles<br>45-150 $\mu\text{m}$ | Non-Microporous | 1ml |
| **T150C** | 100% β-TCP | β-TCP particles<br>150-500 $\mu\text{m}$ | Non-Microporous | 1ml |
| **TCPscaffold** | 100% β-TCP | Disc shape N/A | 60% | $\Phi=5\text{mm}$<br>1.5mm in<br>thickness |
| **PTMC-B45P** | 70/30 (PTMC/BCP) | Disc shape N/A | 70% in<br>macroview | $\Phi=5\text{mm}$<br>5mm in<br>thickness |
| **PTMC-T45P** | 70/30 (PTMC/TCP) | Disc shape N/A | 70% in<br>macroview | $\Phi=5\text{mm}$<br>5mm in<br>thickness |
| **PTMC-T45C** | 70/30 (PTMC/TCP) | Disc shape N/A | 70% in<br>macroview | $\Phi=5\text{mm}$<br>5mm in<br>thickness |
| PTMC | 100% PTMC | Disc shape N/A | 70% in<br>macroview | $\Phi=5\text{mm}$<br>5mm in<br>thickness |

β-TCP particles of two different size ranges, 45-150 μm (T45C) and 150-500 μm (T150C), the β-TCP block (TCPscaffold) were provided by CAM Bioceramics BV, Leiden, the Netherlands. The β-TCP particles were sintered at 1300 °C for 8 hours and did not contain microporous structures. The β-TCP scaffold was also sintered at 1300 °C for 8 hours and had a porosity of 60%.

1,3-trimethylene carbonate of polymerization grade (Boehringer Ingelheim, Germany), stannous octoate (SnOct2, Sigma, USA), and other solvents (Biosolve, the Netherlands) of technical grade were used as received. For the salt leaching procedure to create porous scaffolds, sodium chloride (NaCl) (Merck) crystals were fractioned into a size range of 200-435 μm by being sieved through meshes of the sizes on a Fritsch sieving machine and were stored in a cool, dry place.

The synthesis and characterization of poly(trimethylene carbonate) has been described in details in a previous study(34). The synthesized PTMC polymer was purified by being dissolved in chloroform and precipitated in a five-fold excess of pure ethanol. A salt leaching technique was used to produce porous PTMC scaffolds. The purified PTMC polymer was dissolved into chloroform at a concentration of 5 g/100 ml and then the sieved NaCl crystals were dispersed into the PTMC-chloroform solution by magnetic stirring. The amount of added NaCl crystals took up 70 vol% of the PTMC fraction. Then the mixed PTMC-NaCl dispersion was precipitated in a five-fold excess of pure ethanol and the PTMC-NaCl precipitation was collected and dried under vacuum at room temperature until constant weight. Dried PTMC-NaCl precipitation was compression molded into discs of 5 mm in diameter and 5 mm in thickness at 140°C under a pressure of 3.0 MPa using a Carver model 3851-0 laboratory press (Carver, USA).

The abovementioned BCP particles from Xpand, β-TCP particles from Xpand, and β-TCP particles from CAM, all in the size range of 45 to 150 μm, were mechanically dispersed into the synthesized PTMC polymer in order to create a PTMC-calcium phosphate composite with 30 vol% (equal to 50 wt%) of calcium phosphate particles. The production of PTMC-T45C composite is taken as an example to describe the producing procedure. The T45C particles were dispersed into the PTMC-chloroform solution with a PTMC concentration of 5 g/100 ml by magnetic stirring to form a homogeneous dispersion. The same salt leaching and compression molding technique as how porous PTMC discs were created were applied to create porous PTMC-T45C composite discs. The prepared PTMC-NaCl discs and PTMC-T45C-NaCl discs were then sealed under vacuum and exposed to 25 KGy γ-irradiation from a ^60^Co source (Isotron BV, Ede, the Netherlands) for sterilization. During the sterilization procedure, the PTMC matrices became simultaneously cross-linked(35). To create porous structures, all discs were gently stirred in demineralized water for a period of three days under sterile conditions to wash out the added NaCl crystals. The demineralized water was changed four times a day. Porous discs of PTMC-T45P and PTMC-B45P in the same size were created and sterilized in the same methods as the porous PTMC-T45C composite discs.

### Surgical procedure

All procedures performed on the sheep were in compliance with the international standards on animal welfare and regulations of the Animal Research Committee of University Medical Center Groningen under the project number 5611.

Ten female adult Dutch Texel sheep were included in the *in vivo* study. The abovementioned different calcium phosphate ceramics and PTMC-calcium phosphate composite scaffolds were implanted in the paraspinal muscles in the sheep under general anesthesia (Fig 2). The general anesthesia was induced with 20 mg/kg sodium pentothal and 2.5 ml 50 mg/ml Finadyne and maintained by inhalation of 3% sevoflurane. The implantation sites on the back were marked with non-resorbable sutures (Polypropylene, Ethicon, USA) in muscle fascia. The wounds were closed layer by layer with resorbable sutures (Polyglactin 910, Ethicon, USA). Amoxicillin (15 mg/kg) was administered before surgery and until six days after the operation. Buprenorphine was applied for pain relief before and after surgery.

**Fig 1.**
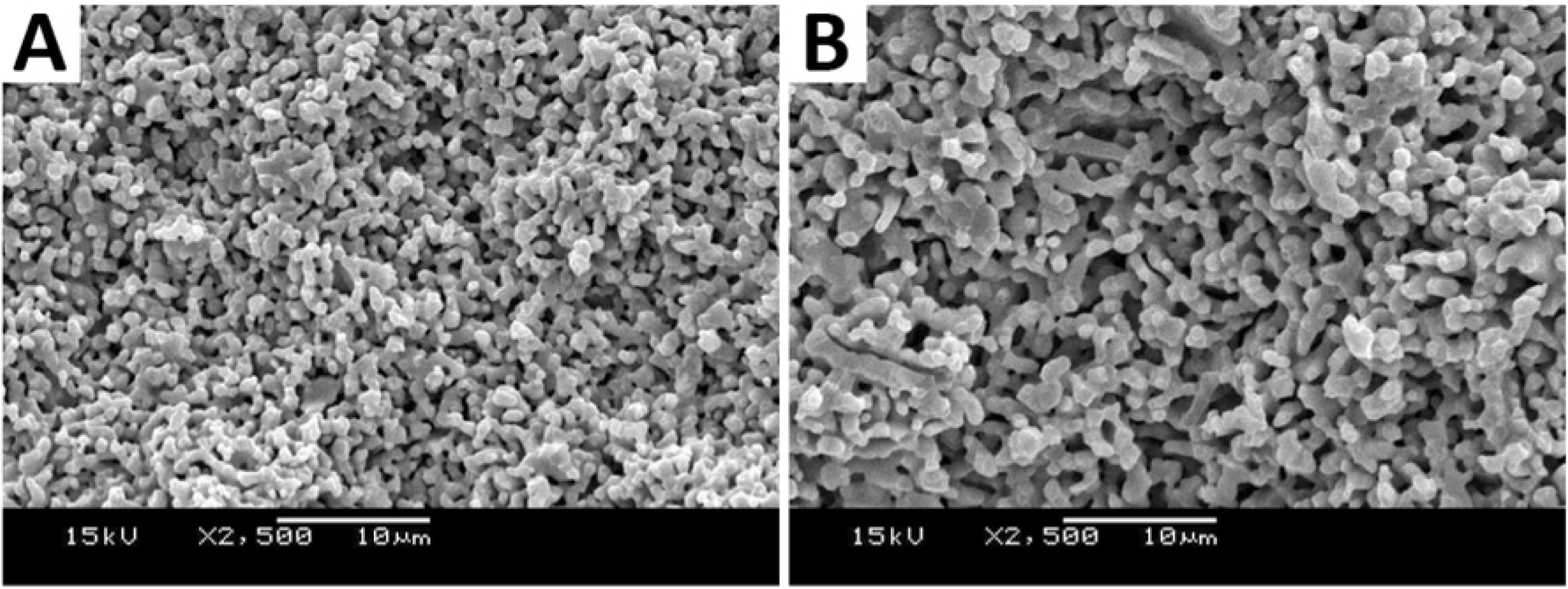
Scanning electron microscope images of β-TCP particles and BCP particles in the size range of 45-150 μm from Xpand. A: β-TCP particles; B: BCP particles. Images are kindly provided by Xpand Biotechnology BV, Bilthoven.

**Fig 2.**
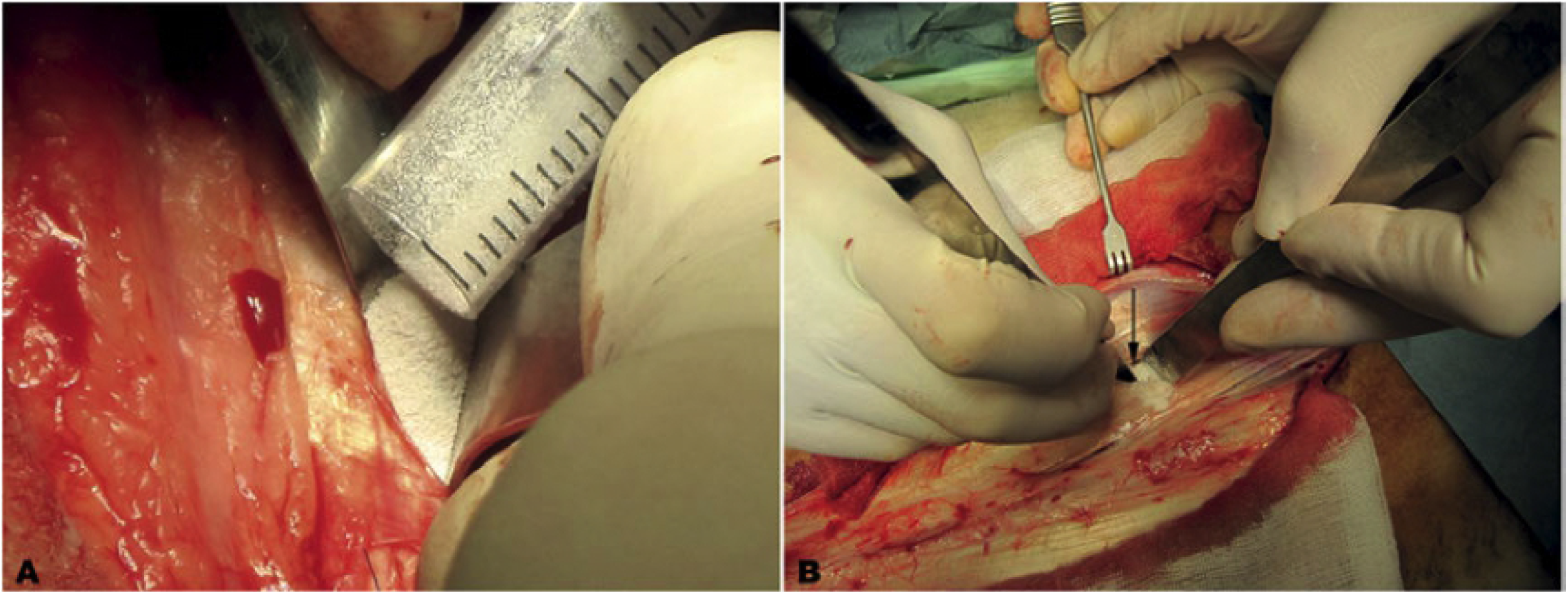
Surgery scheme. A: Placing calcium phosphate particles (B45P, B150P, T45P, T45C, T150C) intramuscularly; B: Placing β-TCP ceramic scaffolds, PTMC-calcium phosphate composite scaffolds (PTMC-B45P, PTMC-T45P, PTMC-T45C), or PTMC porous scaffolds intramuscularly. Different scaffolds are shown by the arrow(↓). Notice that none of the abovementioned biomaterials were fixated in the implantation sites.

Fluorochrome markers were injected to monitor potential new bone formation with time passing by. Table 2 shows the schedule of injections of fluorochromes. The sheep were sacrificed at three months and nine months, five for each time point, respectively, by an injection of overdosed pentobarbital (Organon, the Netherlands). After each termination, muscle tissue containing the implants was retrieved and fixed in a 4% phosphate buffered formalin solution.

**Table 2.** Schedule of fluorochrome injection

|  | Calcein | Xylenol Orange | Oxytetracycline<br>(Engemycin) |
| --- | --- | --- | --- |
| **Supplier** | Sigma, the Netherlands | Sigma, the Netherlands | Mycofarm, the Netherlands |
| **Solvent** | 2% NaHCO<sub>3</sub> | 1% NaHCO<sub>3</sub> | Physiological saline |
| **pH** | 6-8 | 6-8 | 6-8 |
| **Administration** | 10 mg/kg,<br>intravenously | 100 mg/kg,<br>intravenously | 32 mg/kg,<br>intramuscularly |
| **Time of injection**<br>**for 3-month group** | After three weeks | After six weeks | After nine weeks |
| **Time of injection**<br>**for 9-month group** | After 30 weeks | After 33 weeks | After 36 weeks |

### Preparation for histological evaluation

After being rinsed thoroughly with phosphate buffered saline and dehydrated in gradient ethanol solutions, the fixed samples were embedded in methyl methacrylate (LTI, the Netherlands) without being decalcified. Sections of 20-30 μm in thickness for histological evaluation were sawed through the center of the samples parallel to the long axis of sheep dorsal muscle by a modified diamond saw (Leica SP1600, Leica Microsystems, Germany). 1% methylene blue (Sigma-Aldrich, the Netherlands) and 0.3% basic fuchsine (Sigma-Aldrich, the Netherlands) were used to counterstain sections for observations under light microscope. The sections were digitalized by a slide scanner (Dimage Scan Elite 5400 II, Konica Minolta Photo Imaging Inc., Japan) for observations under 25× magnification and by a digital camera on Leica microscope (DFC 420 C, Leica microsystems, Germany) for observations under 200× magnification. Histological evaluation on the sections focused on new bone formation induced by the biomaterials, degradation of the biomaterials, and tissue reactions towards the biomaterials. New bone formation induced by the biomaterials was quantified by their incidence and was scored according to the amount of newly formed bone from zero to three. Zero stood for no new bone formation observed, one stood for limited new bone formation or only mineralization observed, two stood for moderate new bone formation, and three stood for abundant new bone formation. Unstained sections were produced for observations under an epifluorescent confocal laser microscope (Leica TCS SP2, Leica, Germany) to monitor the dynamics of new bone formation, if there was any. The observation was carried out under a 20× oil immersion objective. Calcein with a peak absorption wavelength(abs.) of 500 nm and a peak emission wavelength(em.) of 545 nm displayed green in the images, xylenol orange of 543 nm abs. and 580 nm em. displayed red in the images, and tetracycline of 405 nm abs. and 560 nm em. displayed blue in the images.

### Histomorphometry

Digital images obtained from the slide scanner were used for histomorphometry with Adobe Photoshop CS6. In each image, an overall region of interest (ROI) was determined to contain the implantation site and the newly formed bone, if there was any, and the corresponding pixels were registered. Then newly formed bone, if there was any, was manually selected using ‘Magic Wand Tool” with a tolerance set as ‘50’, and the corresponding pixels were registered. The percentage of newly formed bone was calculated by the following formulation:

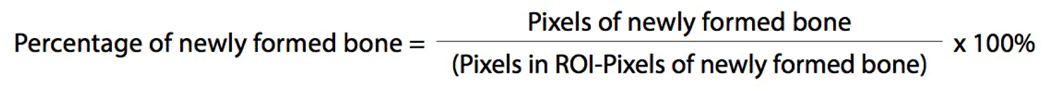

The calculation was performed twice with the researcher blinded from the information of the sections. The mean percentage of newly formed bone and the standard deviation were reported in the results. A Mann-Whitney U test was performed to compare the performances of different materials.

## RESULTS

In a total of 83 implanted samples, results of 11 samples could not be included for the purpose of this study unfortunately. Several reasons contributed to this regrettable loss of samples. First of all, one sheep, which had been healthy, died six months after the surgery and causes for the unexpected death were not revealed by a post mortem autopsy. Only the two BCP samples, namely BCP45P and BCP150P, from the deceased sheep were harvested and included in the nine months follow-up group. The other six samples, namely TCPscaffold, T45C, T150C, PTMC-T45C, PTMC-T45P and T45P, were disposed of with the cadaver. The other sheep did not show signs of infections or other complications during the experiment. Secondly, although non-degradable sutures marked the implantation sites, difficulties existed in relocating and retrieving the implanted biomaterials after time spans of several months, due to the biodegradability of the materials and that the biomaterials were not fixed in the paraspinal muscle tissue. Thirdly, some samples did not contain the region of interest or were not suitable for processing. These last two reasons account for the other five samples (consisting of the following materials: two T45P’s, T45C, PTMC and PTMC-T45C) which could not be included in the study. Fig 3 shows an overview of histological observations of the implanted calcium phosphate bioceramic particles and β-TCP scaffolds under 25× magnification, and Fig 4 shows an overview of histological observations of PTMC-calcium phosphate composite scaffolds and PTMC scaffolds under 25× magnification. Fig 5 shows an overview and detailed images of the implanted TCPscaffold. Fig 6 shows epifluorescent confocal micrographs of TCPscaffold. Fig 7 shows histological observations under 200× magnification concerning new bone formation, degradation of and tissue reactions towards the calcium phosphate bioceramic particles. Table 3 shows the histological evaluation and histomorphometry data on new bone formation induced by different calcium phosphate bioceramic materials.

**Fig 3.**
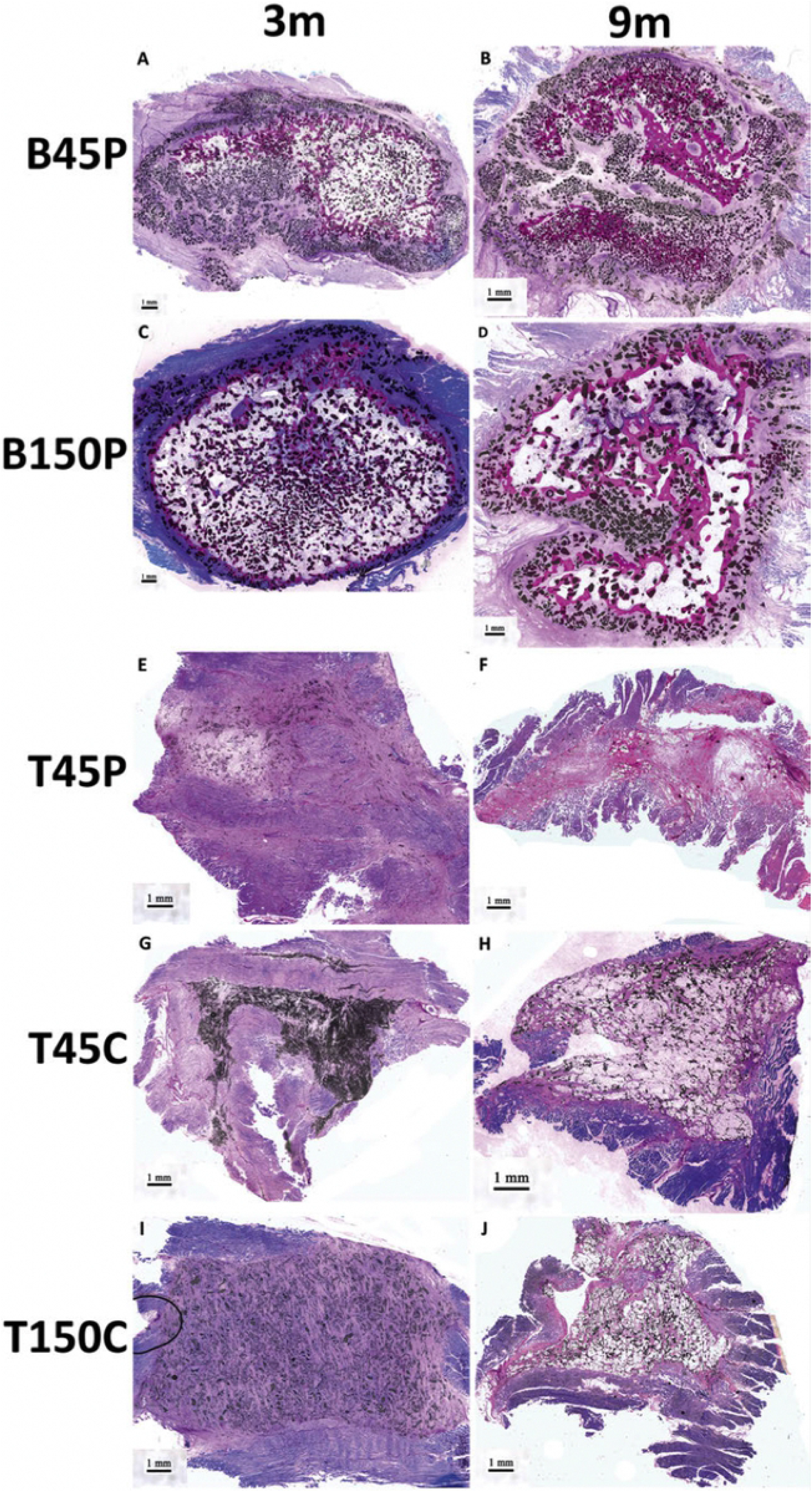
Observations under 25× light microscope of different calcium phosphate bioceramic particles implanted intramuscularly after three months and nine months. Newly formed bone was stained bright red in the sections. Obvious new bone formation was induced by BCP particles of both 45-150 μm and 150-500 μm after both three months and nine months. No new bone formation occurred in the implantation sites of all β-TCP bioceramic particles. Calcium phosphate bioceramic particles were present as black particles in the implantation sites. BCP particles retained their granular shape at both three months and nine months with reduced sizes. β-TCP particles from both sources had disintegrated substantially into clusters of fine particles at three months and the majority of these particles disappeared at nine months. Excessive loose fibrous tissue encapsulated, infiltrated and segregated the implantation sites at both time points. At nine months the implantation sites had been largely replaced by adipose tissue.

**Fig 4.**
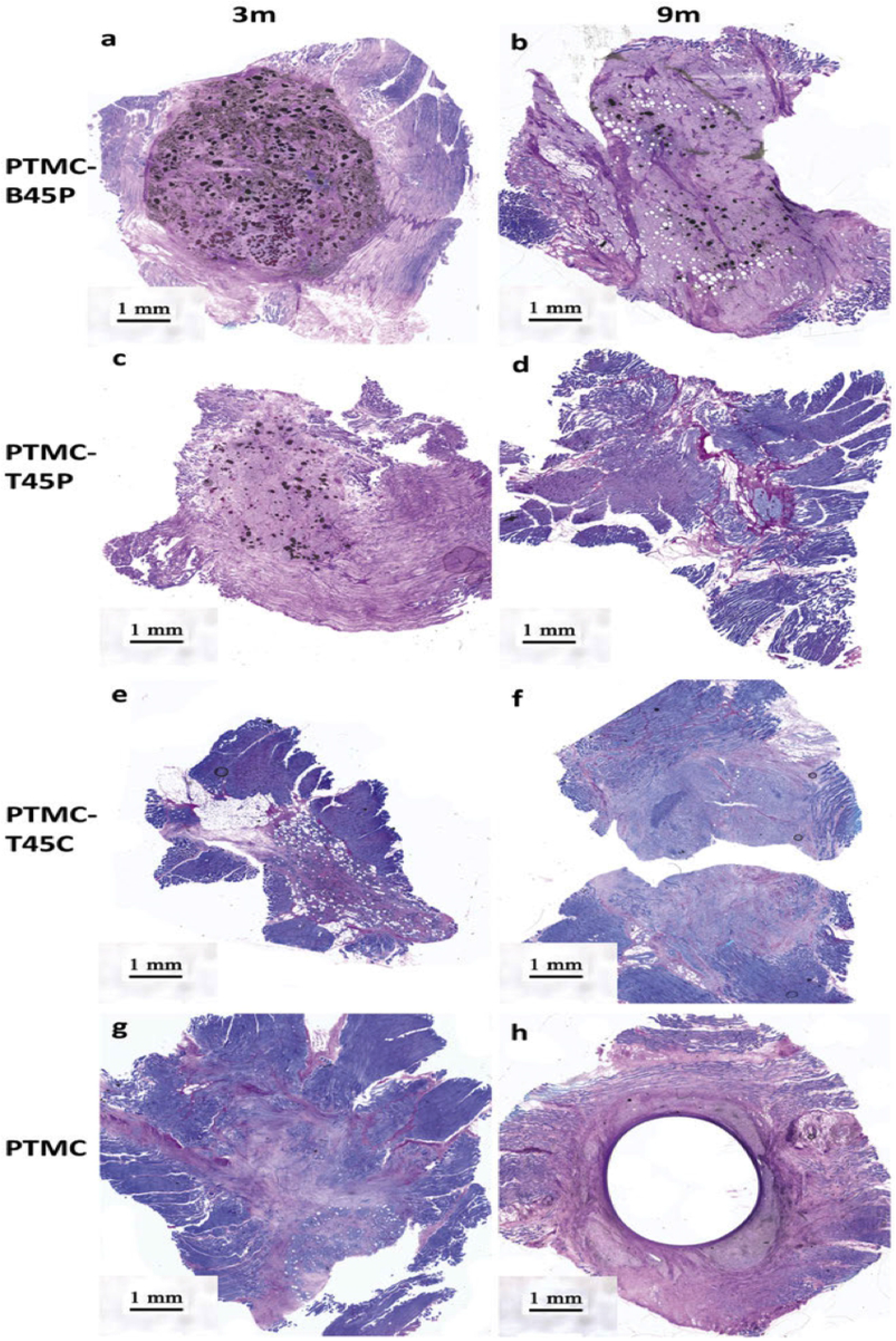
Observations under 25× light microscope of different PTMC-calcium phosphate composite scaffolds and PTMC scaffolds implanted intramuscularly after three months and nine months. No new bone formation was induced by these biomaterials at both time points. Some BCP particles of 45-150 μm remained intact with reduced sizes and the others disintegrated into fine BCP particles at three months. Only a few BCP particles of 45-150 μm were still present in the implantation sites at nine months. Only a few β-TCP particles of 45-150 μm from Xpand were observed in the implantation sites, while β-TCP particles of 45-150 μm from CAM had mostly disappeared at three months. At nine months, β-TCP particles from both sources had disappeared from the implantation sites. PTMC matrices of the PTMC-calcium phosphate composite scaffolds and PTMC scaffolds had been degraded uneventfully into remnants of different sizes at three months and into homogenous small pieces at nine months. Round voids were left in the samples of PTMC scaffolds after nine-month degradation. Excessive loose fibrous tissue encapsulated, infiltrated and segregated the implantations sites at both time points.

**Fig 5.**
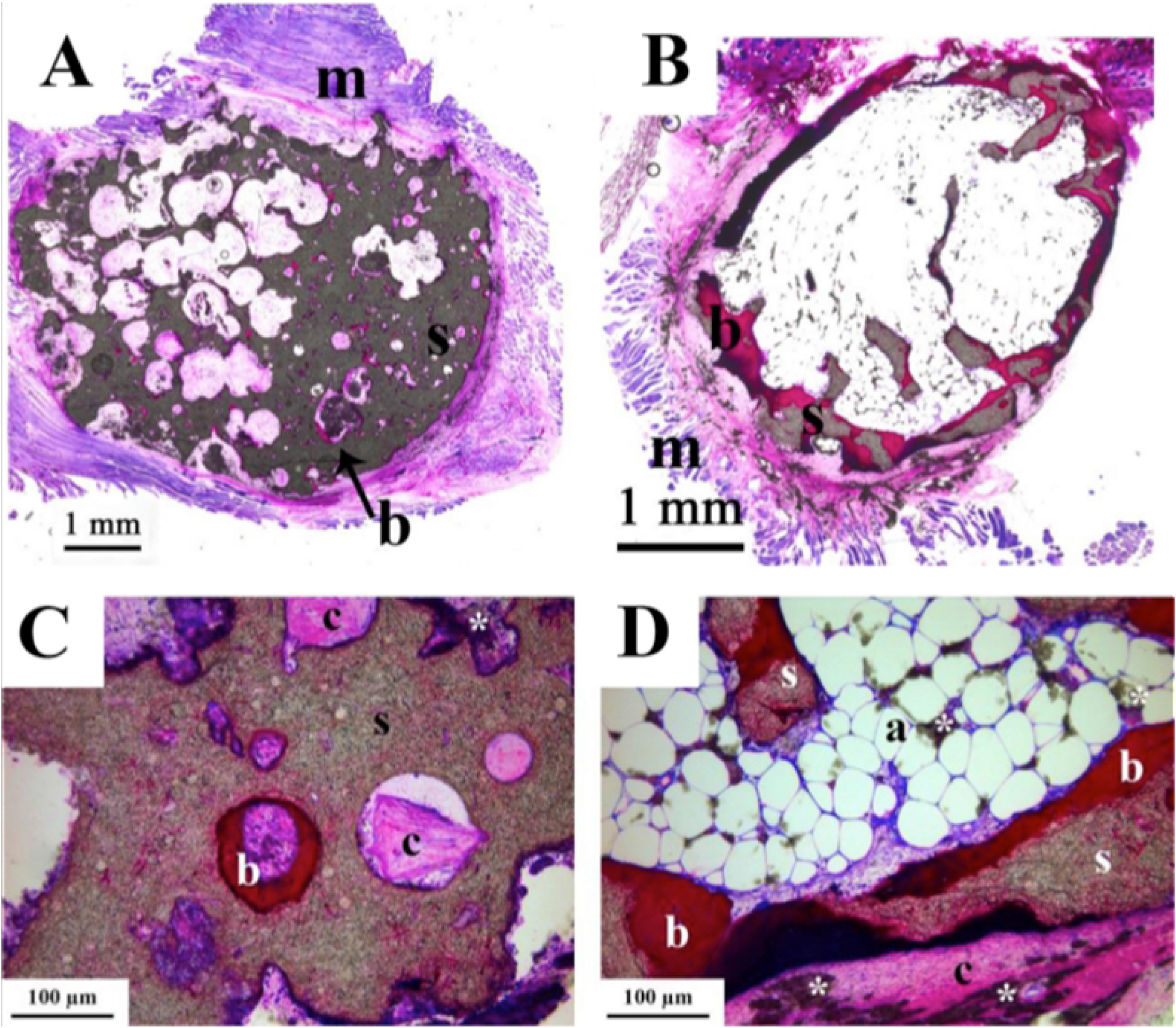
Overview and detailed micrographs of the implanted TCPscaffold. Fig A & C and B & D represent the three and nine month follow-up respectively. Images A and B are overview observations. Images C and D are the corresponding detailed images under 25× light microscope. New bone formation occurred in both follow-up timepoints. Fig A and C show that new bone formation starts in the macropores and still endures when it is in close contact with the scaffold after 9 months (B & D). Newly formed bone (b) can be observed in detail in images C and D. The tissue reactions are visualized in detail in images C & D. After three months, moderate degradation of the TCPscaffold was observed. After nine months, this degradation process continued to the extent in which almost all biomaterial had degraded. a. adipose tissue b: newly formed bone, m: muscle, s: TCPscaffold, c: connective tissue, *: disintegrated TCPscaffold. The scale bar represents 1 mm in images A & B, and 100 μm in images C & D.

**Fig 6.**
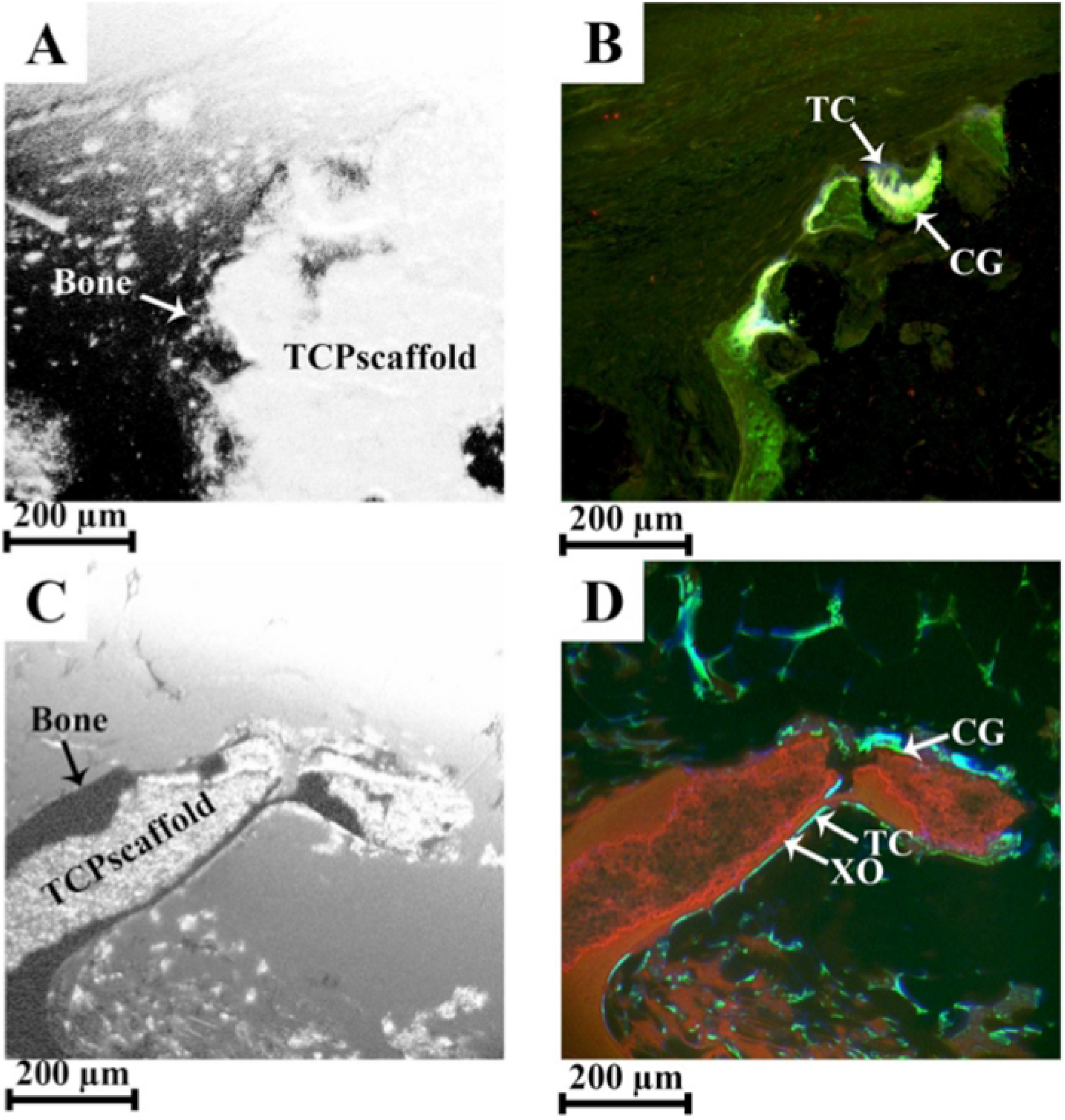
Epifluorescent confocal micrographs of the implanted TCPscaffold. Fig A-B and C-D represent the three and nine month follow-up, respectively. Images A and C are bright field images. Images B and D are the corresponding epifluorescent images. New bone formation started as early as three weeks after implantation, marked by the green colored zones (B) and continued throughout the implantation time. Up until nine months of follow-up, the bone tissue was still expanding and being remodeled since all colors are represented (D). CG: calcein, green; XO: xylenol orange, red; TC: tetracycline, blue. Scale bar represent 200μm.

**Fig 7.**
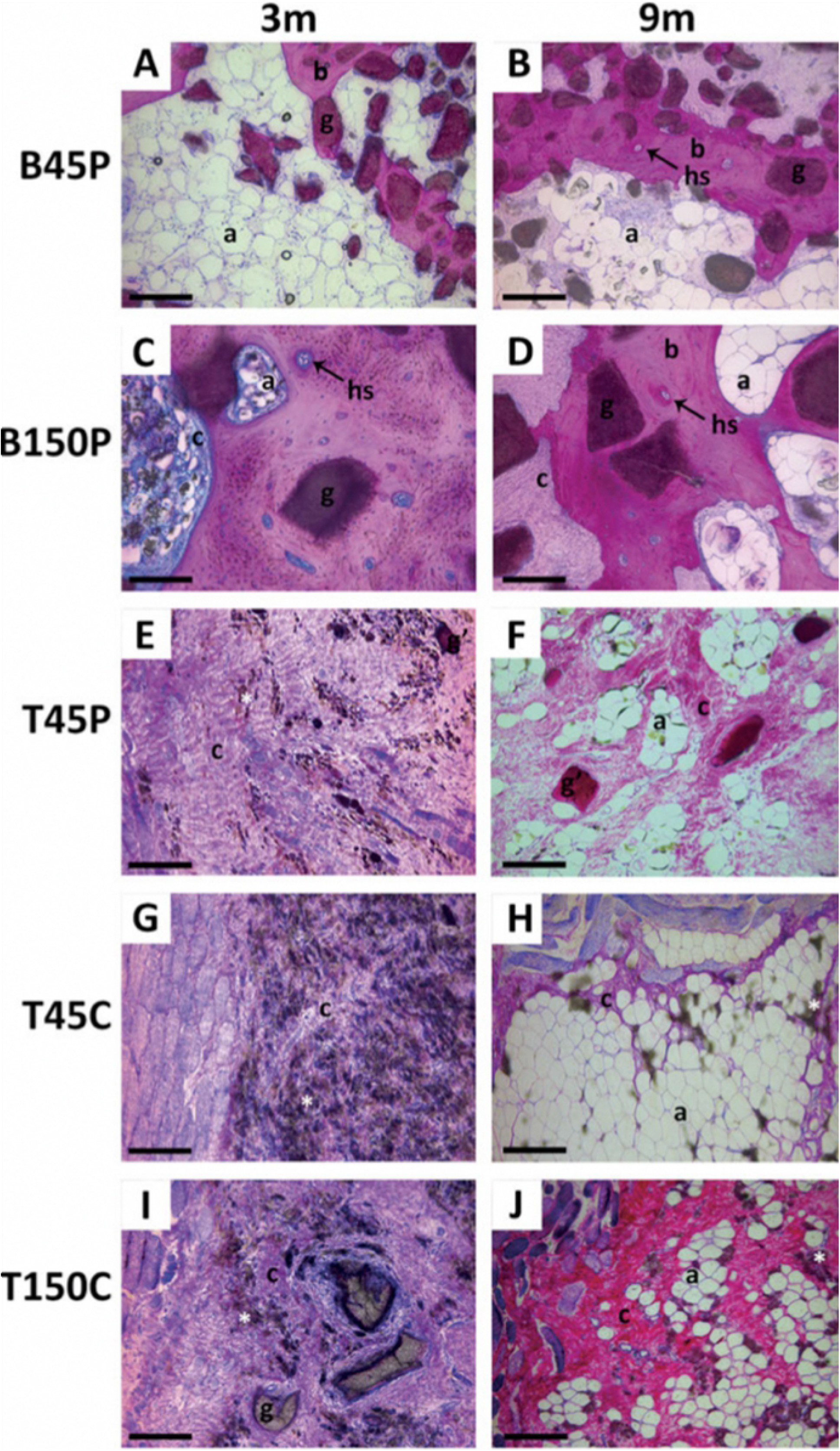
Observations under 200× light microscope of different calcium phosphate bioceramic particles implanted intramuscularly after three months and nine months. New bone induced by BCP particles of both 45-150 μm and 150-500 μm was formed in close contact to the BCP particles at both three months and nine months. b: newly formed bone; HS: Haversian system; g: calcium phosphate particles; g’: red-stained calcium phosphate particles; *: disintegrated calcium phosphate particles; c: fibrous connective tissue; a: adipose tissue. Scale bar represents 50 μm.

**Table 3.**
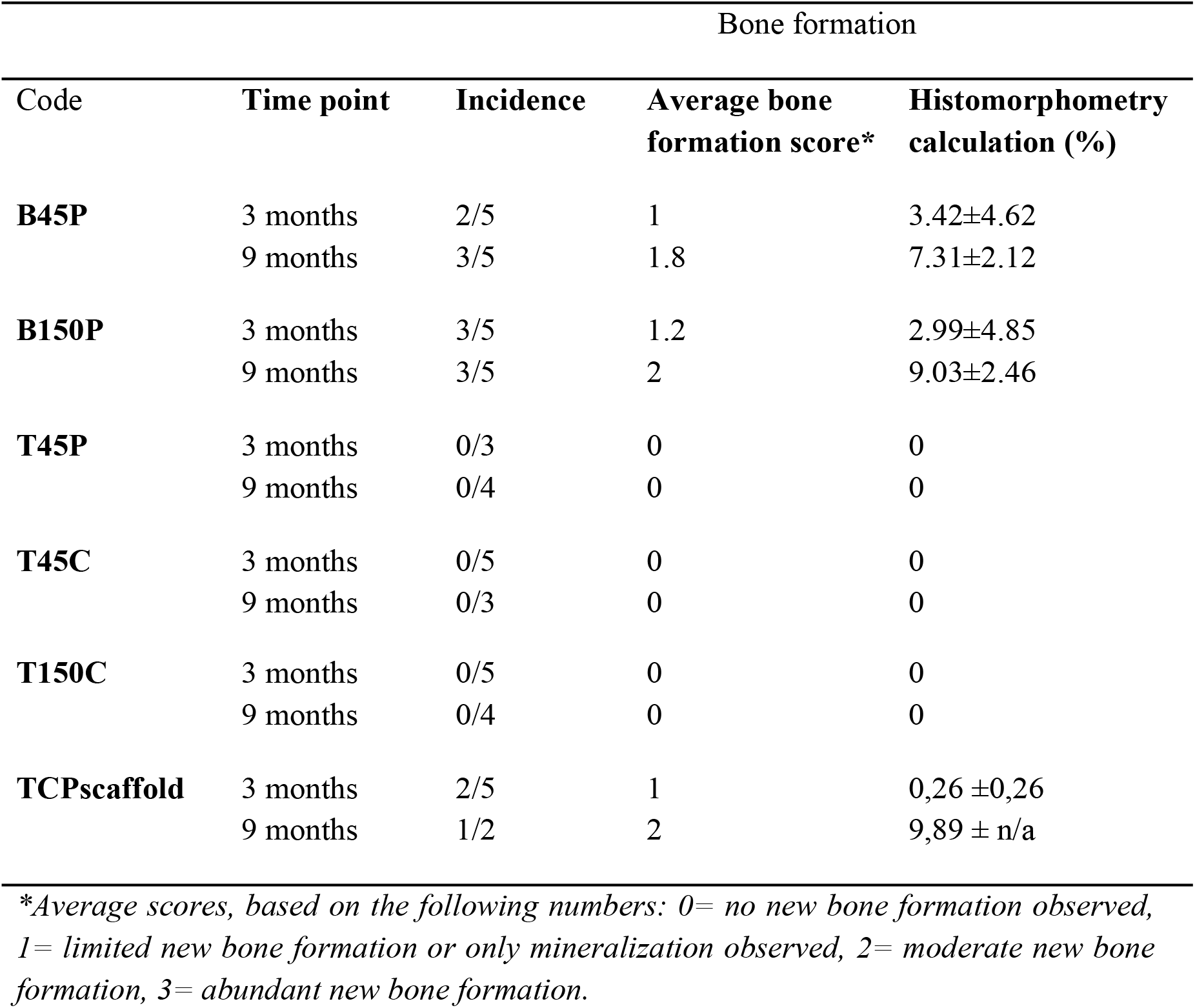
New bone formation induced by different calcium phosphate bioceramic materials

Histological evaluation of the sections was carried out under light microscope to distinguish whether there was new bone formation or not. Abundant new bone formation was seen induced by BCP particles of both size ranges at both three months and nine months. New bone formation was not observed in the groups of different β-TCP particles at neither time points, from neither sources, nor of different size ranges. However, the pure β-TCP (TCPscaffold) was able to induce bone formation in both the three months and the nine months time point, which is an unexpected observation since pure β-TCP ceramics are usually not regarded as being osteoinductive. No new bone formation occurred in the groups of PTMC-calcium phosphate composite scaffolds and PTMC scaffolds, therefore grading of new bone formation and histomorphometry were not carried out for these biomaterials. Induced by BCP particles of both size ranges, new bone formation occurred in close contact to surfaces of the BCP particles (Fig 7 A-D). At three months new bone was formed into continuous patches outlining contours of the implantation sites and scattered small pieces interconnecting remaining BCP particles. At nine months newly formed bone was observed to be of larger amounts and more maturity with a clearly visible Haversian system than at three months. New bone induced by the BCP particles resembled cancellous bone in structure at nine months. As for the amount of newly formed bone, no statistical significance was shown between the BCP particles of both size ranges, regardless of the two time points. Fig 8 shows observations of new bone formation induced by the BCP particles of both size ranges under epifluorescent confocal microscopy. Different colors from fluorochrome markers represented the deposition of newly formed bone in a time order.

**Fig 8.**
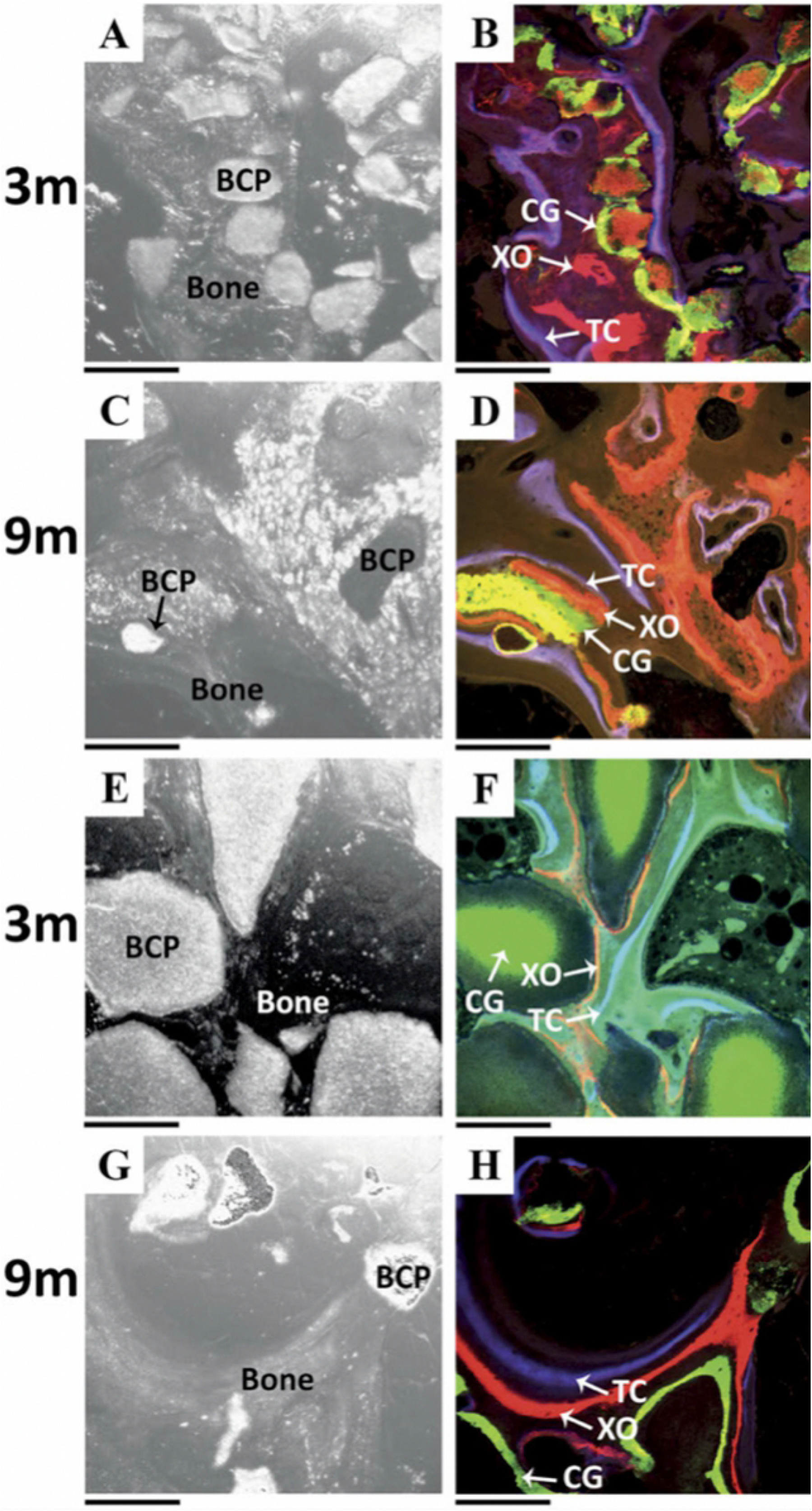
Observations of BCP particles of 45-150μm (A-D) and 150-500 μm (E-H) implanted intramuscularly under epifluorescent confocal microscopy after three months (A, B, E, F) and nine months (C, D, G, H). Images A, C, E and G are observations under light field, while images B, D, F and H are corresponding epifluorescent confocal images. New bone formation started as early as three weeks after implantation and continued through the whole implantation time spans. New bone formation and bone remodeling remained active at nine months. CG: calcein, green; XO: xylenol orange, red; TC: tetracycline, blue. Scale bar represent 200μm.

New bone formation started as early as three weeks after the implantation of the BCP particles and continued until the animals were sacrificed. New bone formation and bone remodeling still remained active at nine months. At three months, disintegration of BCP particles and TCP particles was observed to different extents. Most of the BCP particles of 45-150 μm remained intact with reduced sizes and these intact particles were surrounded by “dust like” disintegrated particles (Fig 3 A). BCP particles of 150-500 μm showed reduced sizes and remained intact. Areas containing the “dust like” disintegrated BCP particles were not seen at the implantation site (Fig 3 C). Compared to BCP particles of both size ranges, β-TCP particles showed advanced disintegration. β-TCP particles of 45-150 μm from Xpand have been obviously disintegrated into clusters of ne TCP particles. Only scarce T45P particles remained intact with a reduced size (Fig 3 E). β-TCP particles from CAM bioceramics of both size ranges were substantially disintegrated and the whole implantation site was filled with TCP particles (Fig 3 G, I). Only few TCP granules of 150-500 μm from CAM bioceramics were seen intact with a reduced size (Fig 3 I).

The macropores of the TCPscaffold (Fig. 5 A & C) facilitated the development of new bone tissue. The contact between the newly formed bone and the surface of the TCPscaffold was excellent and clearly provided mechanical stability since all formed bone was only observed in close proximity of (semi-)intact scaffold-structures. The TCPscaffold was more resistant to degradation than the β-TCP particles. The TCPscaffolds mostly maintained their structure after 3 months, although disintegration and resorption by host tissue took place as well (Fig. 5 A). Of the two TCPscaffold samples, one sample had collapsed completely after 9 months, leaving “dust-like” areas containing TCP particles behind. The other sample, containing mature bone tissue, still contained intact scaffold structures when it was in close contact with the newly formed bone (Fig 5 B & D).

At three months, PTMC matrices had degraded into remnants of different sizes in PTMC-calcium phosphate composite scaffolds and PTMC scaffolds (Fig 4 a, c, e, g and Fig 9 3m). The implantation sites of PTMC-B45P were filled with fine BCP particles that had disintegrated from B45P particles. Some intact B45P particles with reduced sizes were present too. A few remaining B45P particles were stained red on the sections, implying that certain protein absorption occurred on the microporous surfaces of the B45P particles, despite of no new bone formation observed in these samples (Fig 4 a). The T45P particles of the PTMC-T45P composite disintegrated into fine particles and only a few intact T45P particles in reduced size remained. A few red-stained remaining T45P particles were also present at the implantation sites (Fig 4 c). T45C particles had all disintegrated into fine TCP particles and degradation of these fine T45C particles has already progressed much. Very few fine TCP particles scattered at the implantation site of PTMC-T45C (Fig 4 e). At nine months, degradation of all implanted biomaterials had progressed further. Intact BCP particles of both size ranges with reduced sizes were still present at the implantation sites, but the number of these remaining BCP particles had been much reduced compared to at three months. The ‘dust like’ areas consisting of disintegrated BCP particles at the implantation sites of B150P particles were not as prominent as that of B45P particles (Fig 3 B, D). β-TCP particles of both size ranges from both sources had almost completely disintegrated (Fig 3 F, H, J). Only few countable T45P particles with much reduced sizes were seen at the implantation sites (Fig 3 F). No intact β-TCP particles of both size ranges from CAM, but only clusters of fine β-TCP particles were left at the implantation sites (Fig 3 H, J). A few intact BCP particles, although in much reduced sizes, were still present in the implantation site of PTMC-B45P (Fig 4 b). TCP particles of 45-150 μm from both sources went through complete disintegration, and had degradation and disappeared from the implantation sites of PTMC-T45P and PTMC-T45C scaffolds (Fig 4 d, f). PTMC matrices of the PTMC-calcium phosphate composite scaffolds had degraded into homogenous small remnants, which were being processed by macrophages and foreign body giant cells (Fig 9, 9m). Interestingly, degradation of PTMC scaffolds at nine months resulted in discernable round voids surrounded by layers of fibrous tissue (Fig 4 h). This could possibly be explained by a decreased rate of degradation due to a decrease in molecular weight during their degradation procedure(36). The host tissue reacted in a same manner towards all implanted biomaterials (Fig 4, 7). Abundant loose fibrous tissue encapsulated, infiltrated, and segregated the implantation sites at three and nine months. Inflammatory cells, such as macrophages and foreign body giant cells, were presented at the implantation sites, processing disintegrated calcium phosphate particles and degraded PTMC matrices. At nine months, the implantation sites were largely replaced by adipose tissue.

**Fig 9.**
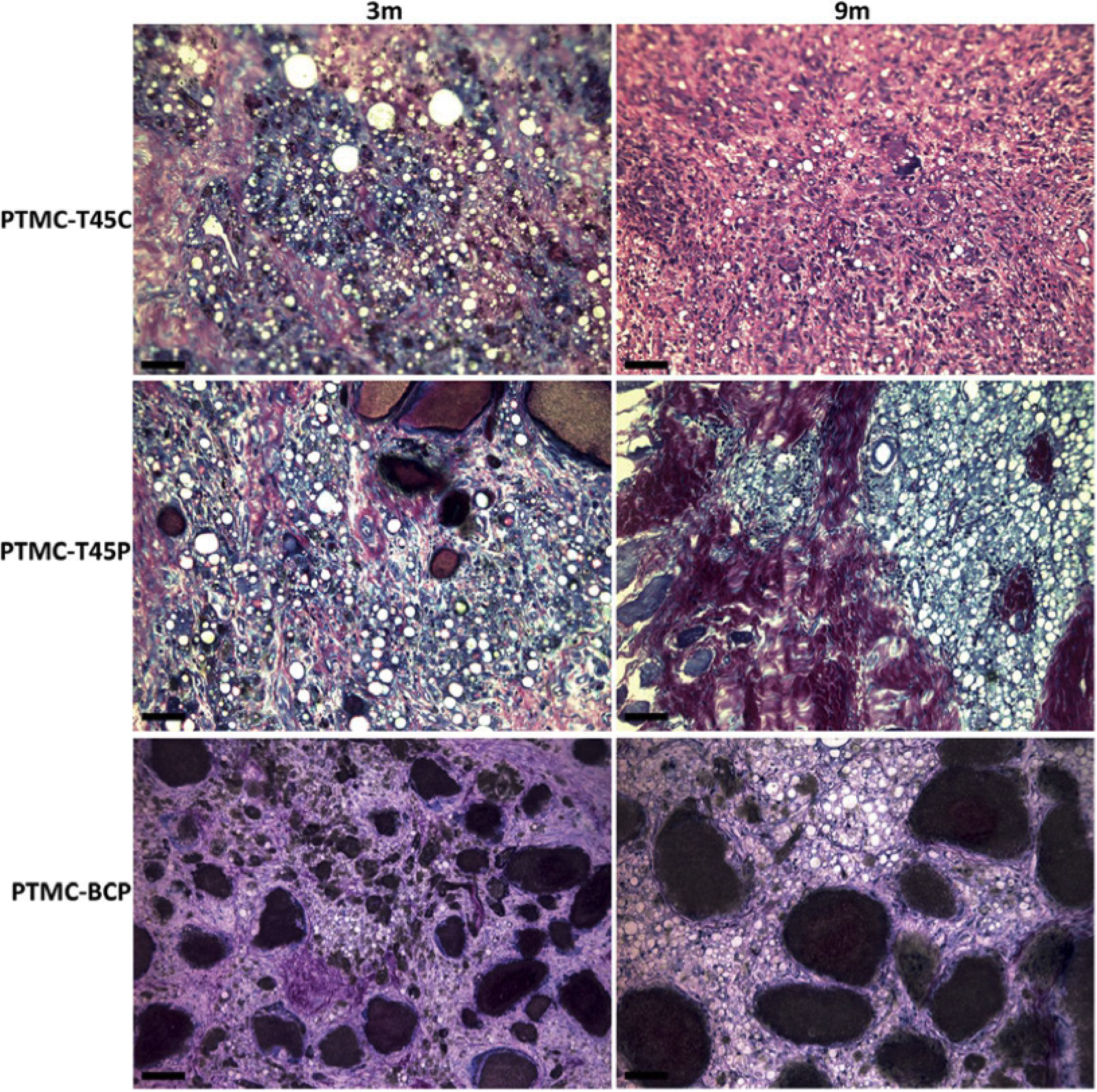
Observations under 200× light microscope of different PTMC-calcium phosphate composite scaffolds implanted intramuscularly after three months and nine months. * PTMC remnants; g: calcium phosphate particles; arrows point at foreign body giant cells. Scale bars represent 50 μm.

## DISCUSSION

This study was carried out to assess osteoinductive properties of different calcium phosphate bioceramic materials and PTMC-calcium phosphate composite scaffolds and to evaluate the *in vivo* degradation characteristics and tissue reactions towards these biomaterials.

Biomaterials with osteoinductive properties have been shown to enhance their regenerative performance in critical orthotopic bone defects(6), thus research on osteoinductive biomaterials becomes an increasingly hot topic. Whether a biomaterial is osteoinductive or not is determined by various factors(7). Crucial determining factors of a ceramic biomaterial being osteoinductive include chemical composition, that is the ratio between β-TCP and HA(28); grain size(29); synthesizing temperature(24); porosity under macroview and microview(15); surface roughness and specific surface area(31). BCP bioceramics have been proven to possess osteoinductive capacities(6, 24) thanks to a balance between the presence of β-TCP, which is more resorbable, and HA, which is more stable and resorption-resistant(20, 27). Implanting β-TCP particles from Xpand, which contained 10% of HA, intramuscularly in sheep led to no new bone formation, indicating the important role of HA in making bioceramics osteoinductive. Mechanical stability of ceramic biomaterials also influences their capacity of inducing new bone formation(26, 30). The β-TCP particles from both sources disintegrated too fast to provide a stable surface for new bone formation. Thus, besides the lack of HA in their composition, the incapability of the β-TCP particles in osteoinduction can also be accounted to a low level of mechanical stability. In our study, BCP particles of two different size ranges were revealed to induce substantial new bone formation in sheep long dorsal muscle. A previous study showed that BCP particles of 45-150 μm were osteoinductive(34). Compared to the previous study, BCP particles of a larger size range showed similar osteoinductive capacity. Mixtures of HA and β-TCP particles (Zimmer, Warsaw, IN) ranging from less than 44μm up to 1000 - 2000μm were seeded with human bone marrow stromal cells and implanted in mice subcutaneously. Mixed HA/TCP particles in a size range of 100 - 500μm led to the most abundant new bone formation, indicating that HA/TCP particles in a size range of 100-500μm provide highly suitable niches for human bone marrow stromal cells to attach, infiltrate, proliferate and differentiate (37), echoing the finding in our study. Besides, BCP particles with a HA/TCP ratio of 60:40 and a size range of 1000 to 2000μm have been shown to induce mature *de novo* bone, which bridges remaining BCP particles and resembles natural cancellous bone after six months of implantation in sheep back muscles(29). Therefore, granule sizes of a wide range influence osteoinductive properties of biomaterials. Sintering temperature strongly influences the microporosity, surface roughness and specific surface area of a ceramic biomaterial. BCP particles and HA particles sintered at relatively lower temperatures possess fine microporous structures within the macropore walls and their specific surface area increases with the decrease of synthesizing temperature(38). A microporous structure in ceramic biomaterials facilitates diffusion of body fluid through the bioceramics and thus provides a better environment for cells to infiltrate and differentiate.(26) BCP particles synthesized at 1150°C enhance the osteogenic differentiation of human multipotent marrow stromal cells(39) and BCP scaffolds produced at 1150°C lead to new bone formation throughout the whole constructs both in iliac wing defects and in intramuscular implantation in goats(6). The BCP particles tested in our study were sintered at 1150°C for 8 hours and resulted in potent osteoinductive capacity. One possible explanation for why TCP particles from CAM Bioceramics sintered at 1300°C did not induce new bone formation intramuscularly, aside from not containing any HA, is the relatively high sintering temperature, which leads to a decrease in their microporosity and specific surface areas. In our study, some BCP particles and TCP particles from Xpand were observed to be stained red in the sections, implying that protein absorption occurred on the rough surfaces of their microporous structure despite that new bone formation happening around those particles. The TCPscaffold, containing 100% β-TCP, was seen to induce bone formation. In as early as three months, areas containing bone tissue were observed throughout the macropores of the scaffold to a very limited extent. After nine months, one sample contained mature, remodeled bone tissue. This was an unexpected observation, since pure β-TCP ceramics are usually not regarded as being osteoinductive. Although reports in which β-TCP scaffolds showed osteoinductive properties. the newly formed bone in these reports could not maintain their structure after a longer time period and did not show bone maturation and remodeling.(39,41) Compared to the non-osteoinductive T45C and T150C particles, the TCPscaffold, which was porous and crafted in a block shape, was able to induce bone tissue. The formation of new bone in the pores of the TCPscaffolds stresses the importance of mechanical stability of biomaterials. All samples containing newly formed bone (BCP granules and TCPscaffold) showed that bone formation only occurred in mechanically stable areas, i.e. areas that contained intact granules or scaffolds. Not only is a stable surface necessary for bone formation to take place within the first months, it is also needed for the bone tissue to maintain its structure. After 9 months all samples containing newly formed bone still had (semi)intact granules or parts of the scaffolds in the nearby area of generated bone tissue. T45P, T150C and T45P particles were too brittle or disintegrated too quickly and did not show any signs of new bone formation. Such an observation can be explained by a lack of mechanical stability. Besides that the TCPscaffold offered a certain degree of mechanical stability, it also provided a porous structure. The porous structure of the scaffolds may have functioned as a ladder for the osteoprogenitor cells to migrate on and facilitated the circulation of body fluids.

None of the PTMC-calcium phosphate composite scaffolds tested in our study induced new bone formation in the sheep long dorsal muscle. Given the 70% porosity in the PTMC-calcium phosphate composite scaffolds, the content of solid material, consisting of calcium phosphate bioceramic particles, was limited to only 30%. Moreover, the ratio of PTMC-calcium phosphate was only 70/30, which means that only 9% of the material consisted of calcium phosphate. Besides a limited amount of calcium-phosphate in the material, the calcium phosphate particles disintegrated fast and provided too low mechanical stability for new bone formation. This could be a reason why the tested PTMC-calcium phosphate composite scaffolds were not osteoinductive. A previous study showed that new bone formation of a limited amount is observed in the center of non-porous PTMC-BCP composite sheets, after the composite sheets are implanted in sheep long dorsal muscles for three months(34). The PTMC-BCP composite sheets contain 30% BCP particles of 45-150μm and are of 1.5 mm thick. Compared to our PTMC-BCP porous scaffolds, the PTMC-BCP sheets contained more BCP particles and were thinner. Since PTMC matrices degrade through surface erosion, such non-porous composite sheets offer more chances for BCP particles to be released into the surrounding tissue and exert their osteoinductivity. That calcium-phosphate containing composites have an osteoinductive potential has been proved by Le Nihouannen et al. They implanted BCP/fibrin composites of 2 to 3 cm^3^ in sheep long dorsal muscles for six months. The implanted BCP/fibrin composites contained BCP particles with a HA/β-TCP ratio of 60/40 and a diameter of 1-2 mm and a fibrin glue of 4 IU. Formation of well-mineralized, matured bone tissue bridging the BCP particles has been confirmed by histology, back scattered electron micrographs and μCT. The number and thickness of the formed bone trabeculae were similar to those in the vertebral body(40).

## CONCLUSIONS

Biphasic calcium phosphate particles of 45-500μm induced abundant *de novo* bone tissue after being implanted in sheep dorsal muscle for three and nine months, and were potently osteoinductive. The fact that new bone formation was seen in the TCPscaffold at both time points suggested that a structure with macropores, a source of calcium phosphate, a favorable surface structure and good mechanical stability could have facilitated the differentiation of osteoprogenitor cells into osteoblasts. Although incorporating β-tricalcium phosphate particles into a matrix of PTMC facilitates implantation, the PTMC matrix impedes the osteoinductive properties of the materials. The porous composite scaffolds composed of PTMC matrices and three different β-tricalcium phosphate particles of 45-150μm induced no new bone formation in sheep dorsal muscle during the implantation periods and showed no osteoinductive capacities. Implantation of different calcium phosphate bioceramic particles of different sizes and these PTMC-calcium phosphate composite scaffolds in sheep long dorsal muscle led to uneventful degradation of the abovementioned biomaterials and provoked no serious tissue reaction. Future studies are needed to determine the optimal composition of composite biomaterials based on PTMC and calcium phosphate to produce osteoinductive composites.

